# Free energy and flexibility analysis of autoinhibited human BRAF

**DOI:** 10.1101/2025.08.04.668576

**Authors:** Jeremy O. B. Tempkin, Fikret Aydin, Sebnem Essiz, Yue Yang, Timothy S. Carpenter, David E. Durrant, Deborah K. Morrison, Helgi I. Ingólfsson, Frederick H. Streitz, Dwight V. Nissely, Felice C. Lightstone, Xiaohua Zhang

**Affiliations:** Physical and Life Sciences Directorate, Lawrence Livermore National Laboratory, Livermore, CA 94550, USA; Molecular Biology and Genetics, Department of Engineering and Natural Sciences, Kadir Has University, Fatih 34083, Istanbul, Turkey; Laboratory of Cell and Developmental Signaling, Center for Cancer Research (CCR), National Cancer Institute (NCI), Frederick, MD 21702, USA; Computing Directorate, Lawrence Livermore National Laboratory, Livermore, CA 94550, USA; NCI RAS Initiative, Cancer Research Technology Program, Frederick National Laboratory for Cancer Research, Leidos Biomedical Research, Inc., Frederick, MD 21702, USA

**Keywords:** . RAF kinase, 14-3-3 protein, molecular dynamics, thermodynamics, conformational change

## Abstract

The RAF serine/threonine protein kinases function as direct effectors of RAS in the intracellular transmission of extracellular growth signals, and they are key targets for drug discovery given the high incidence of oncogenic mutations in RAF and other components of this signaling pathway. In its inactive state, RAF is held in an autoinhibited conformation in the cytosol through a combination of intramolecular interactions and binding to a regulatory 14-3-3 protein dimer. Activation of RAF is initiated by its interaction with membrane-localized, GTP-bound RAS, which induces conformational changes that release RAF from its autoinhibited state. However, the molecular mechanisms governing RAF activation remain incomplete, largely due to the challenges in experimentally capturing intermediate conformational states in this process. To address this gap, we developed a comprehensive all-atom model of BRAF based on existing cryo-EM structures. Using this model, we performed extensive molecular dynamics simulations to evaluate the stability and free energy landscape of autoinhibited BRAF in solution. Our analysis reveals conformational flexibility within the autoinhibited complex, suggesting that this dynamic behavior may play a role in facilitating BRAF activation upon engagement with membrane-bound RAS.

## INTRODUCTION

The RAF family of kinases, comprising ARAF, BRAF, and CRAF, occupies a central position in the MAPK signaling cascade, transmitting extracellular signals from receptor tyrosine kinases to downstream effectors such as ERK (1). Among these isoforms, BRAF has gathered considerable attention due to its frequent mutation and dysregulation in various cancers, including melanoma, colorectal cancer, and thyroid cancer (2).

Structurally, BRAF contains several distinct domains, including an N-terminal RAS-binding domain (RBD), a cysteine-rich domain (CRD), and a C-terminal kinase domain (KD). A 14-3-3 dimer (14-3-3_2_) binds to phosphorylated serines 365 and 729 (referred to here as 14-3-3 (S365) and 14-3-3 (S729), respectively), stabilizing the autoinhibited conformation and occluding both the dimer interface region of the KD and the membrane binding region of the CRD. Upon growth factor engagement of receptor tyrosine kinases (RTKs), RAS becomes activated and recruits BRAF to the plasma membrane via high-affinity interactions with the RBD. Once recruited, BRAF undergoes structural modifications. Release of 14-3-3 from the serine 365 site and the subsequent dephosphorylation of this site by the SHOC2-MRAS-PP1 complex enables CRD to interact with membrane lipids and RAS to facilitates RAF dimerization for full KD activation (1, 2).

Additionally, in the cryo-EM structure of the autoinhibited BRAF complex (PDB 7MFD) (3–5), key ionic bond-forming residues on the RBD that mediate high affinity RAS binding are largely exposed. However, structural overlay of RAS onto the complex reveals steric clashes between RAS and 14-3-3, indicating that conformational rearrangements of the RBD, 14-3-3, or both are likely required to permit full RAS engagement. Despite these insights, the precise mechanistic details governing the transition from the autoinhibited BRAF-14-3-3_2_ complex to signaling-competent conformations, remain poorly understood. Furthermore, the exact sequence of structural events leading to RAF activation is still unclear. A deeper mechanistic understanding of these transitions and structural states could reveal novel targets for therapeutic intervention.

To investigate the intrinsic flexibility of RAF and the initial conformational changes required to relieve autoinhibition upon RAS binding, we constructed a comprehensive all-atom (AA) model of the autoinhibited BRAF:14-3-3_2_ complex in solution, based on a previously reported cryo-EM structure (3). Our model encompasses BRAF residues 156 to 738, including two large intrinsically disordered regions that connect the cysteine-rich domain (CRD) to the kinase domain (KD) and surround the pS365 binding site for the 14-3-3 dimer (14-3-3_2_). Using this model, we performed extensive AA molecular dynamics (MD) simulations to characterize the conformational landscape around the fully autoinhibited state in solution. We applied essential dynamics analysis and state-of-the-art free energy sampling methods to quantify both the lowest vibrational models near the autoinhibited state and the relative free energy barriers associated with specific inter-domain interactions known to be implicated in BRAF activation. Our results reveal that the lowest-frequency motions correspond to a hinge-like movement that displaces the KD away from the CRD and the 14-3-3_2_ dimer interface. Further, the KD interface with the CRD is considerably more flexible than the interface between the 14-3-32 dimer and the CRD, having a free energy barrier to disruption comparable to that of the RBD:14-3-3 interface. These findings suggest that the motions of the KD domain may constitute a critical step in the BRAF activation process by facilitating the disruption of autoinhibitory interactions to promote RAF dimerization and activation.

## METHODS

### Building the complete autoinhibited BRAF model

We constructed our model of the autoinhibited BRAF:14-3-3_2_ complex based on the recent cryo-EM structure of the BRAF:14-3-3_2_:MEK complex (PDB: 7MFD) (3). To build the model for MD simulations, several additions and modifications were made as described below. First, MEK was removed from the complex to reduce system complexity and computational cost (further discussed in the Results and Discussion section). ATP and its coordinating Mg^2+^ ion were then placed using Molecular Operating Environment (MOE) (6) (v2022.02) with PDB 6U2G (chain B) serving as the template (Supplementary Information Figure S1). Adjustments to the contacting amino acids were made to reflect the ATP and Mg^2+^ binding in PDB 6U2G, followed by a short energy minimization to relax the ATP-Mg complex. We also restored a missing residue at the N-terminus of the 14-3-3 proteins, near the interface of the 14-3-3 dimer, as well as two residues in the loop region (E70, G71). The C-terminus was capped without reconstructing the 15 missing residues (D231 to N245) as this region is distance from the interface. To complete the coordination of one of the zinc-binding sites in the CRD, we added a water molecule to fulfill the zinc coordination requirements and adjusted its placement to ensure proper coordination of the H235 residue (Figure S2). The remaining missing residues (listed in Table S1) were added using the MOE protein structure preparation model using default parameters. We added phosphorylation sites to the model at S365, S446 and S729 as these sites are phosphorylated in the quiescent state of BRAF (4, 7). We elected to use a charge of −1 for each phosphorylation site.

Finally, we added the missing intrinsically disordered regions that connect the C-terminus of the CRD to the N-terminus of the KD (Figures 1 and S3), which were unresolved in the cryo-EM structure 7MFD. This linker is divided into two distinct segments, named here loop I (residues 274 to 359) and loop II (residues 371 to 448). Loop I connects the C-terminus of the CRD to the S365 regulatory site while loop II connects the C-terminal side of the S365 site to the N-terminal side of the KD (Figure 1). To incorporate these disordered regions into our model for MD simulations, we generated a randomized ensemble of initial loop placements using the following procedure. First, we modeled an ensemble of putative loop conformers into the cryo-EM structure using the MOE Loop Modeler with default de-novo loop building parameters. For each loop region, we generated 100 independent loop conformers. From these, we manually selected a representative subset based on conformational diversity and energy score, yielding 9 conformers for loop I and three conformers for loop II. We then combined all possible loops I and II pairs, resulting in an initial library of 27 distinct structural models (Figures 1E and S3).

**FIGURE 1.**
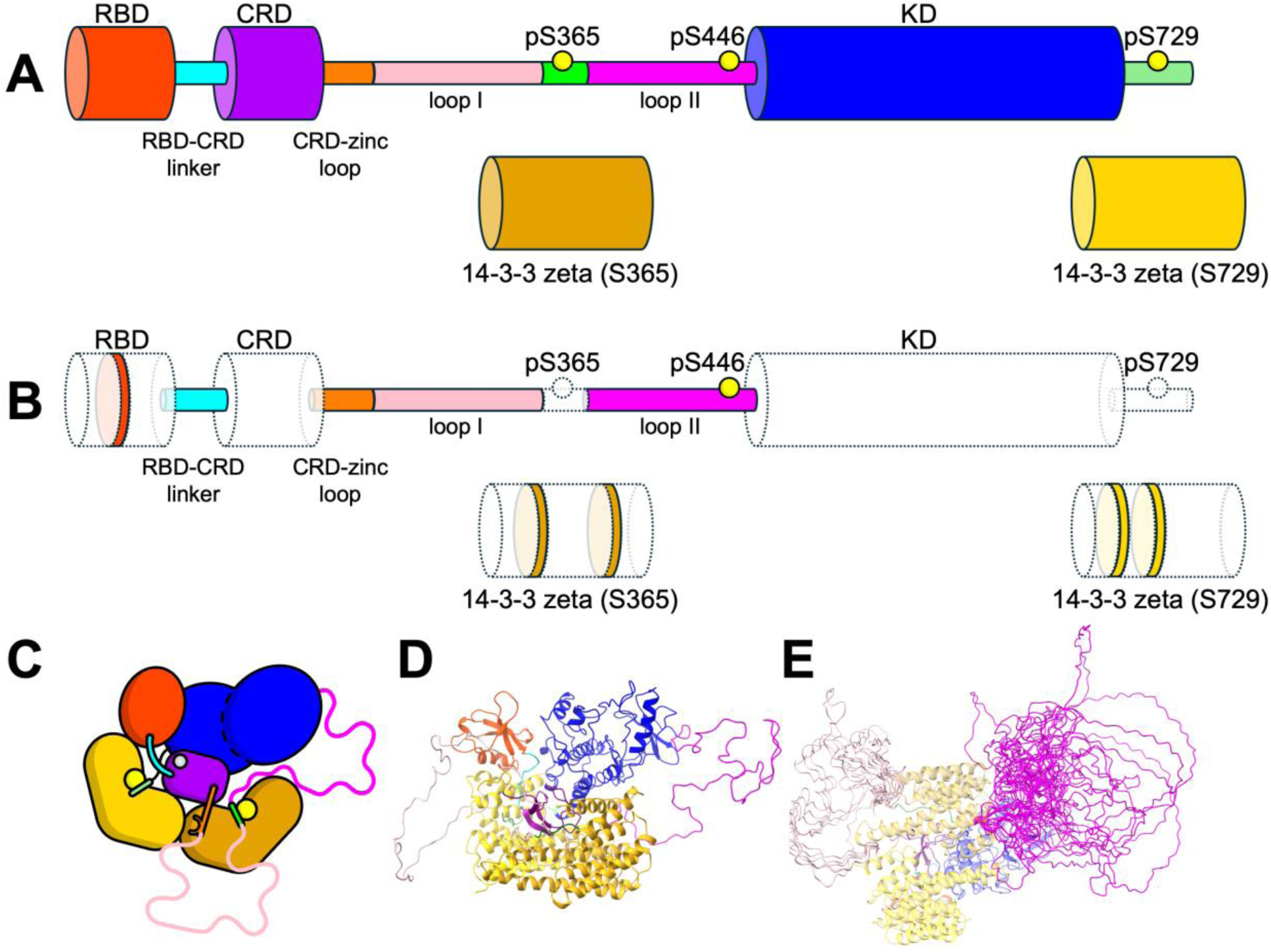
Atomistic model of the complete BRAF-14-3-3_2_ complex in solution. A) Cartoon illustration of the BRAF-14-3-3_2_ complex. Thicker tubes indicate globular domains (RBD, CRD, KD, and both the 14-3-3 protomers in red, purple, blue, gold and yellow, respectively), and thinner tubes indicate linker regions or disordered sections (in cyan, orange, pink, green and magenta). Phosphorylation sites at residues S365, S446 and S729 included in our model are indicated by the yellow circles. B) Cartoon of the BRAF-14-3-3_2_ complex illustrating which sections of the structure were modeled into the original cryo-EM structure (PDB 7MFD). Domains and residues that are resolved in the cryo-EM structure are shaded (white fill with dashed lines) and sections that were modeled in are colored. C) Cartoon of the autoinhibited BRAF-14-3-3_2_ complex showing what domains are in contact using the canonical coloring for each domain. D) Example of the initial structure of the BRAF-14-3-3_2_ complex after initial simulation relaxation using unbiased MD. E) Illustration of the library of initial disordered loops I and II built with MOE loop modeling procedure. Loops I (pink) and II (magenta) are show superimposed on a single BRAF-14-3-3_2_ complex structure aligned to the 14-3-3_2_. View in panel E is from 90-degree rotation from view in panel D.

For each initial conformer, we initialized and simulated 5 independent unbiased MD simulations using the AMBER18 package (8–10). All input files were generated using CHARMM-GUI (11, 12) with standard settings. All MD simulations were then performed using the CHARMM36m force field (13) with a 4 fs timestep (enabled by hydrogen mass repartitioning (14)) unless otherwise noted. The TIP3P (15) water model was used, with water constraints applied via SETTLE (16) and all other hydrogens constrained using SHAKE (17). These unbiased AA simulations were run with mixed-precision (SPFP (18)) AMBER18, following initial topology construction and minimization with GROMACS (v2019.6) (19) and format conversion with the gromber tool of ParmEd from AmberTools 19 (20). Each individual MD simulation within this set was simulated for a minimum of 500 ns each. This procedure generated a library of 135 independent simulations containing a total of 73.2 μs of simulation data.

### Essential dynamics analysis of unbiased simulations

As described in the model-building section, we generated 27 distinct structural models of the autoinhibited BRAF complex, representing all possible combinations of the loop I and loop II conformations. We used a library of 135000 frames from the unbiased MD simulations by selecting the final 1000 frames (last 100 ns of 500 ns) from each of the 135 unbiased MD simulations described above. For the five replicas of each different loop combination, we checked the radius of gyration (RoG) and in the last 100 ns a compact structure had been reached for all simulations. We computed the covariance matrix of the Cα positions and diagonalized the covariance matrix to predict the dominant motions and their corresponding magnitudes around equilibrium structure of the BRAF:14-3-3_2_ complex. Eigenvectors and eigenvalues of the covariance matrix were calculated using the ProDy software (21).

Loops I and II structures are excluded from the principal component analysis (PCA). In the first step, each conformation in the trajectory was superimposed into the initial structure. In the second step, variance-covariance matrix was constructed by calculating the deviations of positions from the average structure. The elements of the covariance matrix, C was calculated as:

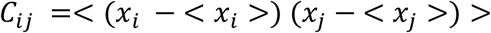

where 𝑥_𝑖_ and 𝑥_𝑖_ are the coordinates of residues i and j, and the brackets denote the ensemble average taken over our conformer library. The covariance matrix, C, can be decomposed as:

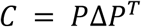

where the eigenvectors, *P*, represent the principal components (PCs) and the eigenvalues are the elements of the diagonal matrix, Δ. Each eigenvalue is directly proportional to the variance it captures in its corresponding PC. The eigenvectors represent the intrinsic collective motions of the protein and the corresponding eigenvalues represent the magnitudes of these motions. To observe the collective motions and its magnitudes, normalized correlations of the systems was calculated as:

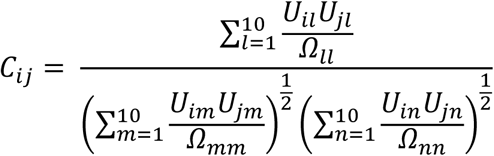

where U is the matrix of eigenvectors, and Ω is the diagonal matrix of eigenvalues. The cross-correlations maps indicate correlation of motion between different parts of the protein.

### 1D umbrella sampling

We manually selected 6 inter-domain distances to serve as coordinates for computing free energy profiles using umbrella sampling (22, 23): 1) RBD to KD, 2) RBD to 14-3-3 (S729), 3) CRD to 14-3-3 (S729), 4) KD to 14-3-3 (S729), 5) KD to 14-3-3 (S365), and 6) CRD to KD. Each inter-domain distance is defined as the distance between the center of mass (COM) of every Cα atom in each domain. All 1D US simulations were performed using OpenMM (v8.0.0) (24). The temperature was maintained at 310K using Langevin dynamics as implemented in the OpenMM package. For each coordinate, we divided the inter-domain distances with windows spaced evenly over the intervals: 1) 2 to 8 nm, 2) 1.4 to 7.4 nm, 3) 1 to 7 nm, 4) 3 to 9 nm, 5) 3 to 9 nm, and 6) 2 to 8 nm, respectively. For the RBD to KD coordinate, we divided the target distance interval into 31 windows, while for all other distance we used 61 windows. We restrained each simulation using a harmonic potential with 1000 KJ/mol*nm force constant and minimum placed at each window center, expect for the RBD to KD distance for which we used a force constant of 500 KJ/mol*nm. We equilibrate the system in each umbrella sampling window by selecting the final frame from one of the unbiased MD simulations described above and restraining the system at the initial inter-domain distance. We then progressively shift the center of the restraint from the initial value to the target position using 20 stages, starting from the initial position and ending at the target restraint position for each window. Each stage was simulated for 2.5 ns with a fixed window center before moving to the next restraint location, totaling 50 ns to equilibrate the system in each window. All production simulations were then simulated with a fixed harmonic restraint for 12 hours on Frontier or Lassen supercomputing systems, resulting in ∼20-40 ns of simulation time per window. This procedure was repeated for each of the 27 loop conformers and across all inter-domain distances, resulting in an aggregate 501.8 μs of total simulation time. All analysis code and simulation setup were implemented in Python and Bash scripting languages.

We computed the 1D potentials of mean force (PMF) using the Weighted Histogram Analysis method (WHAM) (22, 25) implemented in the software from the Grossfield lab (v2.1.0) (26). For each window, we extracted the time series of COM distances and subsampled those trajectories using the timeseries module of the pymbar software (27, 28) into a set of statistically independent samples. We then aggregated the subsampled data across simulations initialized from different starting loop conformations and used this combined dataset as input to the WHAM software. For each pulling direction we computed the PMF over 40 evenly spaced bins spanning between 1) 2.5 to 9.0 nm for RBD to KD, 2) 1.5 to 8 nm for the RBD to 14-3-3 (S729), 3) 1.5 to 7.0 nm for the CRD to 14-3-3 (S729), 4) 3.5 to 7.5 nm for 14-3-3 (S729) to KD, 5) 3.5 to 9.0 nm for 14-3-3 (S365) to KD, and 6) 2.5 to 9.0 nm for CRD to KD. We estimated the variance of each free energy profile using the bootstrapping feature of the WHAM code using 100 independent MC trials.

### 2D metadynamics simulations

Metadynamics is an enhanced sampling technique designed to investigate physical phenomena that occur at time and length scales beyond the reach of standard MD simulations (29, 30). This method enhances the sampling of rare events by adding a history-dependent bias energy into the system’s Hamiltonian. This bias energy takes the form of a Gaussian potential, as defined by:

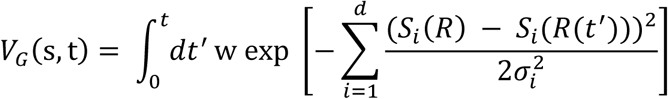

where 𝜎 represents the width of the Gaussian for the i^th^ collective variable (CV), 𝑤 is the height of the Gaussian, and 𝑆_i_(𝑅(𝑡′)) denotes the value of the 𝑖^th^ CV at time 𝑡′.

By continually adding these Gaussian potentials, metadynamics modifies the energy landscape of the system, effectively accelerating the sampling of rare events. This bias discourages the system from revisiting previously explored regions in the CV phase space, promoting more thorough exploration and ideally leading to ergodic and diffusive motion along the chosen CVs.

As the simulation progresses and the bias potentials accumulate, the system is driven to explore new configurations. The sum of the negative bias energies added during the simulation provides an estimate of the underlying free energy profile along the selected CVs. This free energy profile is a powerful tool for understanding the thermodynamics and kinetics of the system, offering insights into the mechanisms of complex processes that are otherwise difficult to capture with standard MD simulations.

Here metadynamics simulations spanning ∼90 ns (15 replicas) were carried out using an approach similar to the one-way sampling approach described previously (31) to determine the free energy surfaces for the 14-3-3 interaction with BRAF when RBD is bound or unbound. The total simulation time for each case is 1.35 µs. The simulations were started after the initial equilibration using the same parameters described above. All simulations were performed using the GROMACS 2019.6 package patched with the PLUMED 2.6.6 code (32–34).

The CVs were selected to describe the motion of the 14-3-3 protomer with S729 site. The use of two CVs has been shown to provide more stable results than the use of one CV (35). The first CV (d_1_COM) is the distance between the COM of the first half of 14-3-3 (S729) (the region closer to CRD) and CRD, whereas the second CV (d_2_COM) is the distance between the COM of the second half of 14-3-3 (S729) (the region closer to RBD) and CRD. The Gaussian bias energy is deposited every 1000 steps with a height of 0.25 kJ/mol and widths of 0.08 nm for the CVs that describe d_1_COM and d_2_COM.

## RESULTS AND DISCUSSION

### Structural heterogeneity of the disordered regions loops I and II

Loops I and II, totaling approximately 150 residues in length (Figure 1), are disordered in solution and are likely unresolved in the cryo-EM structure (3), with the exception of the ∼10 residue section around the serine 365 site. To assess the impact of loop placement on the conformational landscape of autoinhibited BRAF, we generated a large ensemble of initial configurations through randomized position of loop I and II, followed by relaxation and sampling of the conformational landscape via unbiased MD simulations (Figures 1 and S3). Although the initial positions of the loops are qualitatively similar, the relaxed ensemble sampled from the unbiased MD simulation shows that the loops are flexible, sampling the full space around the BRAF complex, with no apparent high affinity for other domains within the autoinhibited complex. Plotting the average density of loop occupations at low, medium, and high threshold values shows that the loops sample a broad area around the complex and only have high occupancy at the positions where the loops are stably bound to the complex (Figure S3). Further, the relaxation of loops I and II show that the ensemble is less extended than in the initial placement (Figure S3).

To confirm proper equilibration of the loop configurations, we plotted the radius of gyration (RoG) and RMSD of the disordered loops (Figure S4) starting from their initial positions (Figures 1 and S3). All trajectories appear stable at RMSD about 25-40 Å for loop I and 10-25 Å for loop II after ∼200 ns of simulation. Outside of the relaxation of loops I and II, no other significant conformational changes in the folded domains were observed in the unbiased simulations, indicating that the overall model is energetically stable around the autoinhibited state.

### Principal components in the essential dynamics analysis reveals the inherent flexibility of the RBD and kinase domain (KD) in the autoinhibited solution structure

To investigate the conformational flexibility of the BRAF-14-3-3_2_ complex in solution, we computed the essential dynamics around the autoinhibited state over a large ensemble of structures derived from our library of unbiased MD simulations (Figure 2). We focused on the first two principal component analysis (PCA) modes, which together represent ∼36% of the observed fluctuations around the autoinhibited state. In the first mode (Figure 2A, C), the RBD and KD open in a hinge-like motion around the point where the CRD interfaces with the 14-3-3 dimer. Simultaneously, the CRD moves outward from this hinge region. In the second mode (Figure 2B, D), the RBD and KD exhibit a twisting motion, again centered at the CRD:14-3-3 interface, resulting in the RBD and KD rotating away from each other. In both modes, the RBD and KD are the most mobile regions of the complex, while 14-3-3_2_ dimer remains comparatively rigid.

**FIGURE 2.**
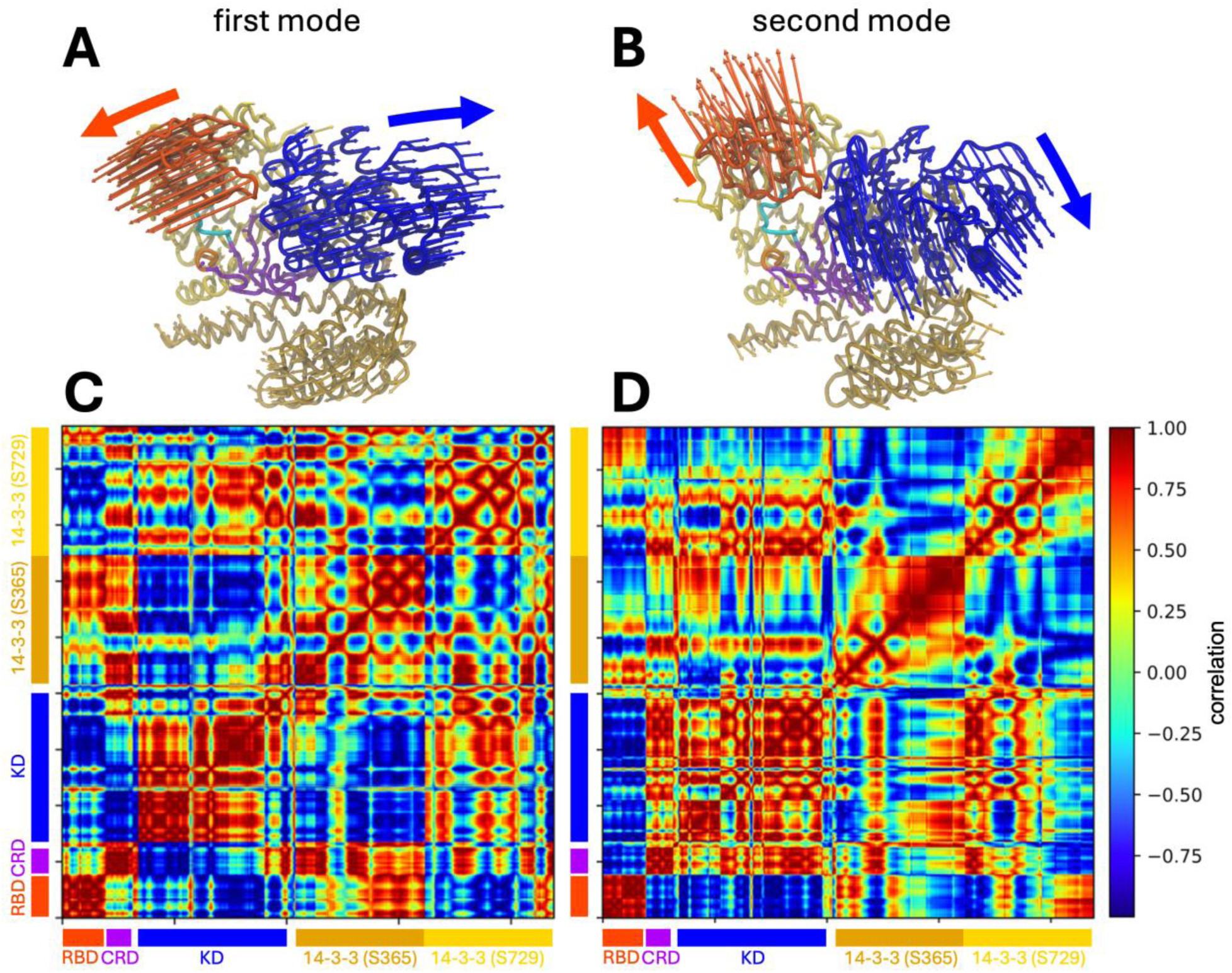
PCA analysis of the autoinhibited BRAF:14-3-32 complex. The proteins/protein-domains are shown in RBD-red, CRD-purple, KD-blue, 14-3-3 (S365)-gold, 14-3-3 (S729)-yellow. A) Shows the 1st PCA mode visualization and (C) the cross-correlation map of the mode. B) and D) show the same for the 2nd PCA mode. The colored arrows indicate the primary motions of each mode.

To further characterize these motions, we computed the normalized cross correlation between domain movements (Figure 2C, D). Interestingly, the CRD displays opposing behavior between the first and second modes: in the first mode, the CRD motion is anti-correlated with that of the KD, whereas in the second mode, it moves in concert with KD. These results indicate that equilibrium fluctuations around the autoinhibited state are primarily driven by motions involving the RBD and KD.

Finally, to assess whether the conformational flexibility around the autoinhibited state is related to the conformational changes required for the KD to transition between the monomeric and dimeric states, we compared these motions to the structural differences observed between the monomeric. (PDB: 7MFD) and dimeric (PDB: 7MFF) cryo-EM structures of BRAF (3). We observed a qualitative similarity between the structural transformation required to convert the dimeric to the monomeric state and the motion described by the second PCA mode, suggesting that this mode may facilitate the KD dimer-to-monomer transition (Figure S5).

Overlaying RAS onto the autoinhibited BRAF structure reveals a steric clash between RAS and the 14-3-3 dimer when high-affinity ionic bonds are formed between the RBD and RAS. This observation suggests that the conformational flexibility within the complex is likely required to accommodate full RAS engagement. Consistent with this expectation, the RBD exhibited relatively high mobility in our unbiased MD simulations. However, a more unexpected finding was the substantial motion observed in the KD, despite the complex being in an autoinhibited state. PCA analysis of the simulation trajectories revealed that the dominant motions around the autoinhibited conformation involve both the RBD and KD, whereas the CRD and the 14-3-3 dimer remain comparatively stable. These results raise the intriguing possibility that KD mobility may play a functional role early in the RAF activation process, potentially priming the complex for conformational rearrangement upon RAS engagement.

### The free energy landscape of BRAF in solution shows that the relative flexibility of the KD is similar to the RBD

As the BRAF:14-3-3_2_ complex reorients and begins to disassemble, the conformational changes leading to BRAF activation are most likely to proceed through the lowest free energy path connecting the fully autoinhibited complex state to the monomeric pre-dimer state. In the earliest stages of this process, the inter-domain interactions observed in the autoinhibited state are likely to remain intact. Thus, these transitions are expected to be dominated by the intrinsic inter-domain free energies surrounding the autoinhibited state. Therefore, mapping this landscape, particularly the free energy associated with inter-domain motion in solution, may provide valuable insights into the likely sequence of conformational changes that occur following RAS engagement, shedding light on the earlies events in the BRAF activation process.

To further characterize the stability of the BRAF autoinhibited state in solution, we mapped the free energy landscape of the complete autoinhibited BRAF:14-3-3_2_ complex in solution using extensive one-dimensional umbrella sampling simulations (23). We selected 6 inter-domain coordinates that span critical interactions between the RBD, CRD, KD and both 14-3-3 protomers (Figure 3): 1) RBD to KD, 2) RBD to 14-3-3 (S729), 3) CRD to 14-3-3 (S729), 4) 14-3-3 (S729) to KD, 5) 14-3-3 (S365) to KD, and 6) CRD to KD. Each coordinate was defined as the distance between the centers of mass (COM) of all Cα atoms within the respective domains. These coordinates were chosen to capture the key inter-domain motions hypothesized to initiate the activation process (Figure 3). The first two coordinates describe the positioning of the RBD with respect to the 14-3-3 (S729) protomer and KD. Coordinates 3 to 6 describe the positioning of the KD and the CRD with respect to the 14-3-3 dimer. To account for the conformational heterogeneity of the disordered loops, we repeated the umbrella sampling procedure for each of the 27 loop conformers, resulting in a combined simulation time of over half a millisecond. While loops I and II are dynamic (Figure S3), metastable interactions between the loops and structured domains can bias the free energy estimates if simulations are too short or limited in number. Our ensemble-based approach mitigates this risk and better captures the broader thermodynamic behavior of the autoinhibited complex.

**FIGURE 3.**
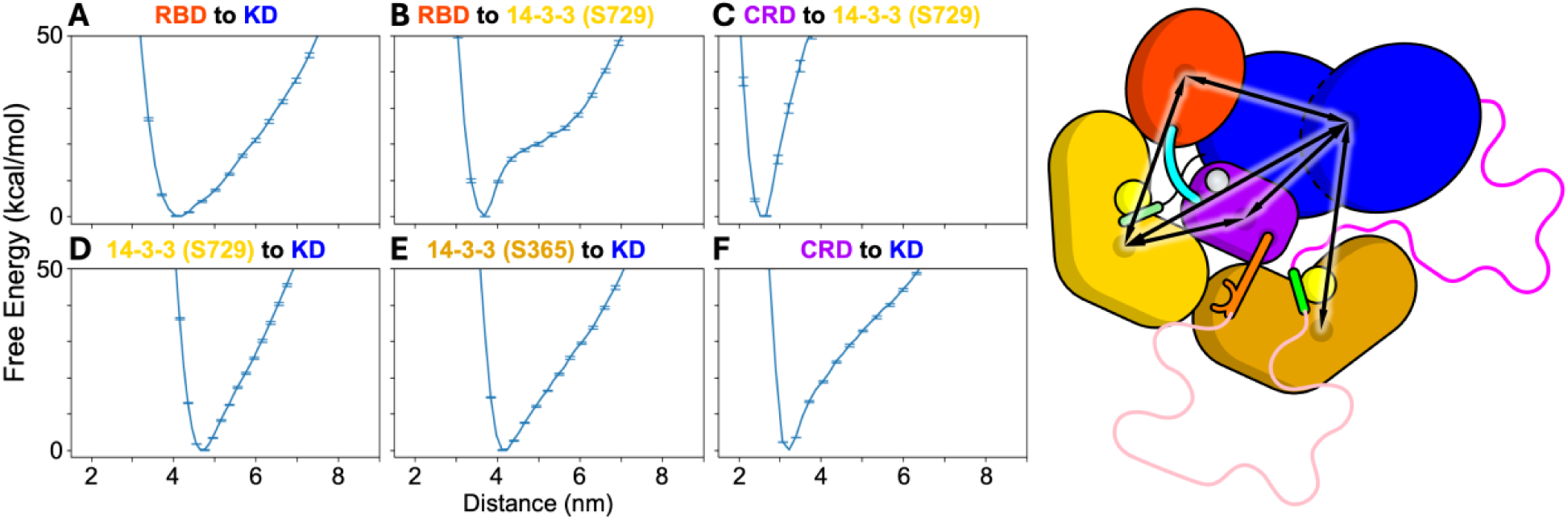
Free energy surfaces of inter-domain distances in the autoinhibited BRAF:14-3-32 complex computed by 1D umbrella sampling. Inter-domain distances are computed as the distance between the COM of all Ca atoms in each domain for the following pairs of domains: A) RBD to KD, B) RBD to 14-3-3 (S729), C) CRD to 14-3-3 (S729), D) KD to 14-3-3 (S729), F) KD to 14-3-3 (S365), and G) KD to CRD. The free energies are estimated by grouping samples from similar windows across each of the 27 independent 1D umbrella sampling simulations which were started from each loop structure. The estimate of the free energies at each window are computed using the WHAM. Error bars are estimated from MC bootstrapping procedure provided in the WHAM software using 100 iterations, indicating the 95% confidence interval around the mean. A cartoon of the autoinhibited BRAF:14-3-32 complex is shown indicating all the inter-domain distances (right).

The 1D free energy profiles indicate that the KD and the RBD exhibit the greatest conformational flexibility around the autoinhibited state, consistent with the normal mode analysis of the complex (Figures 2 and 3). The free energy profiles along the RBD to 14-3-3 (S729) coordinate show that the RBD is bound to the 14-3-3 (S729) protomer with a ∼20 kcal/mol energy barrier (Figure 3). Between ∼4.3 and ∼5.8 nm the RBD dissociates from its native interface with 14-3-3 (S729) but remains tethered to the CRD via a short 6-residue disordered linker. The rise in free energy over this distance range corresponds to the progression extension of this linker. Beyond ∼5.8 nm, the linker between the RBD and CRD is fully extended, and further separation of the domains requires disruption of interactions between the CRD and the 14-3-3 dimer. Further, the free energy profile for the RBD to KD coordinate shows that the RBD position with respect to the KD is relatively stable, likely due to the persistent binding of the RBD to 14-3-3 (S729) in these simulations.

The free energy profiles for the pulling directions between the CRD and both 14-3-3 protomers show that these interactions are stably maintained in the autoinhibited state, with a high energetic cost to disrupt them. Further, the coordinates between the KD and both 14-3-3 protomers are similarly stable. However the energy gradient for separating the KD from the 14-3-3_2_:CRD complex is less steep than that observed for the CRD:14-3-3_2_ interactions, suggesting a somewhat more flexible interface. In contrast, the KD to CRD direction shows that the KD is bound to CRD with an energy barrier of ∼20 kcal/mol, which is comparable to the free energy required to dissociate the RD form the 14-3-3 (S729) protomer. Beyond a separation of ∼4 nm, further displacement of the KD requires extension of the intrinsically disordered loops I and II.

### Disengaging the RBD from 14-3-3 (S729) increases the flexibility of the CRD to 14-3-3_2_ interface

For membrane-bound RAS to fully engage the RBD of BRAF, the native interface between the RBD and the 14-3-3 (S729) protomer must be at least partially disrupted (3). To investigate how disruption of the RBD–14-3-3 (S729) interface affects the conformational free energy landscape of BRAF, we performed a series of two-dimensional (2D) metadynamics simulations starting from two distinct conformational states: one in which the RBD forms a native interface with 14-3-3 (S729) protomer (“RBD-engaged” state), and one in which this interface is broken (“RBD-disengaged” state) (Figure 4).

**FIGURE 4.**
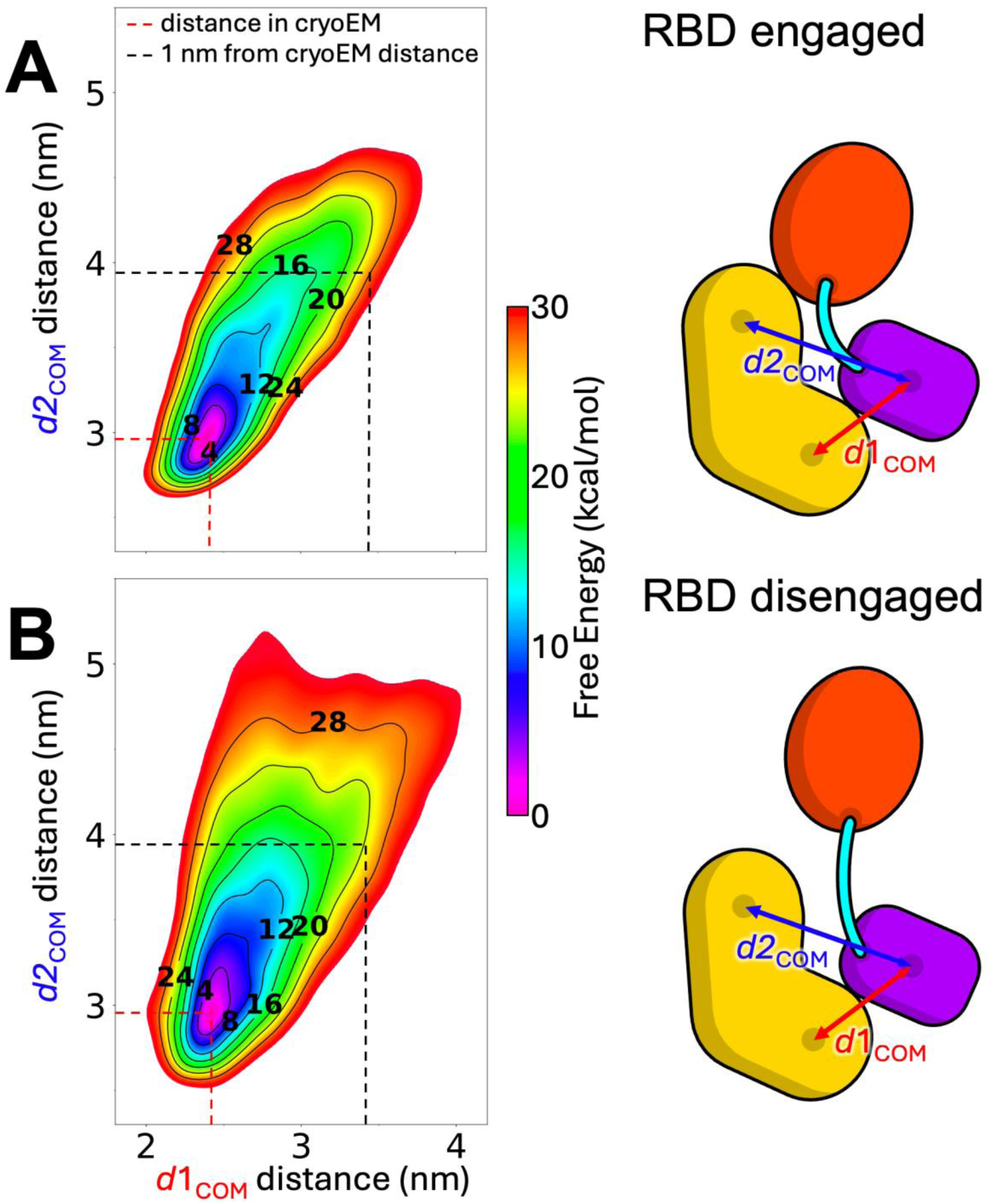
Binding of RAS influences the stability of the 14-3-3 (S729)-CRD interface. 2D potential of mean force (in kcal/mol) obtained by averaging multiple independent metadynamics simulations for understanding the conformational mobility of 14-3-3 (S729)-CRD interface when RBD is bound (A) or unbound (B) to 14-3-3 (S729). The collective variables (CVs) d1COM and d2COM indicate the COM distances between the first half of 14-3-3 (S729) (the region closer to CRD) and CRD, and between the COM of the second half of 14-3-3 (S729) (the region closer to RBD) and CRD, respectively.

The resulting 2D potential of mean force (PMF) plots (Figure 4) capture the motion of the 14-3-3 (S729) protomer relative to the CRD along two collective variables (CVs). As described in the Methods section, CV1 represents the center-of-mass (COM) distance between the CRD and the N-terminal half of the 14-3-3 (S729) protomer (proximal to the CRD), while CV2 tracks the COM distance between the CRD and the C-terminal half of the same protomer (closer to the RBD). These CVs were chosen to bias the system toward conformations in which the 14-3-3 protomer moves away from the CRD, allowing exploration of higher-energy, partially dissociated states. Although the interactions at the CRD–14-3-3 interface were disrupted during sampling, the 14-3-3 protomer remained tethered at the phosphorylated S729 residue.

The free energy landscapes reveal that the motion of the 14-3-3 (S729) protomer is more restricted in the RBD-engaged state than in the RBD-disengaged state (Figure 4A vs. 4B). In the engaged configuration, large conformational rearrangements are energetically disfavored, confining the 14-3-3 protomer to a narrower region of conformational space. This rigidity is likely stabilized by a network of hydrogen bonds formed at the RBD–14-3-3 interface, as described in previous structural studies (3).

In contrast, the RBD-disengaged state permits greater conformational freedom of the 14-3-3 protomer, highlighting how RBD displacement may be a critical step in destabilizing the autoinhibited conformation. These findings suggest that RAS binding, by trapping the RBD in a disengaged state, could facilitate the conformational transitions necessary for BRAF activation.

## CONCLUSION

To characterize the conformational flexibility of autoinhibited BRAF, we constructed a comprehensive all-atom model of the BRAF:14-3-3₂ complex and investigated its dynamics through extensive molecular dynamics simulations and free energy calculations. Our analysis of the essential dynamics and inter-domain free energy landscapes confirms that the BRAF:14-3-3₂ complex is stable in its autoinhibited state.

Among the components of the complex, only the disordered loops I and II exhibit a significantly dynamic behavior (Figures 1 and S3). Quantification of inter-domain motions revealed a hinge-like, low-frequency movement of both the RBD and KD away from the CRD and the remainder of the complex (Figure 2). These findings are consistent with the lower free energy barriers associated with displacing the RBD and KD (Figure 3). The relatively high mobility of the RBD is expected, as the RAS-binding interface on the RBD is partially occluded in the autoinhibited structure (3), necessitating RBD displacement for full RAS engagement. Moreover, RAS binding has been experimentally shown to enhance BRAF activation. Notably, disengagement of the RBD from the 14-3-3 (S729) protomer also increases the conformational mobility of the 14-3-3 (S729) protomer relative to the CRD (Figure 4), thereby reducing energetic barriers for subsequent steps in the release of autoinhibition.

Surprisingly, the KD also demonstrates a relatively low barrier for displacement and a similar low-frequency, hinge-like motion away from the CRD and 14-3-3_2_. This finding suggests that KD movement may play an active role in the early stages of BRAF activation. This mechanism is supported by recent experimental studies showing that stabilizing the KD—through either MEK binding (36) or BRAF inhibitors (37) —can suppress BRAF activation by reinforcing the autoinhibited conformation.

Taken together, our findings support a model in which BRAF autoinhibition is maintained by a network of stabilizing inter-domain interactions. While stable on short timescales, these interactions may fluctuate or weaken over longer periods, particularly in the presence of RAS or other modulatory factors. Such external influences could either promote re-engagement of the autoinhibited state or facilitate transitions toward activation, providing mechanistic insights into how RAF regulation is dynamically controlled at the molecular level.

## ASSOCIATED CONTENT

### Supporting Information

This article contains supporting information. A Supporting Information .pdf is provided containing figures and more detailed information on the system set up, system stability and PCA analysis.

## AUTHOR INFORMATION

### Author Contributions

The manuscript was written through contributions of all authors. All authors have given approval to the final version of the manuscript. J.O.B.T performed the US simulations and analysis, F.A. performed the metadynamics simulations and analysis, X.Z. and F.A. coordinated building the protein model, X.Z. ran the unbiased simulations, and S.E. did the PCA analysis. ‡These authors contributed equally. The authors declare that they have no conflicts of interest with the contents of this article.

## ACKNOWLEDGMENT

This work was performed under the auspices of the US Department of Energy (DOE) by Lawrence Livermore National Laboratory under Contract DE-AC52-07NA27344; and under the auspices of the National Cancer Institute (NCI) by Frederick National Laboratory for Cancer Research (FNLCR) under Contract 75N91019D00024. This research used resources of the Oak Ridge Leadership Computing Facility (OLCF), which is a DOE Office of Science User Facility supported under Contract DE-AC05-00OR22725 and Livermore Computing (LC). For computing time, the authors thank the Livermore Institutional Grand Challenge for time on Lassen and the Advanced Scientific Computing Research Leadership Computing Challenge (ALCC) for time on Summit. For computing support, the authors thank LC and OLCF staff. Release: LLNL-JRNL-2007515.

## ABBREVIATIONS

MD: molecular dynamics
AA: all-atom
PCA: principal component analysis
PDB: protein data bank
HPC: high performance computing
cryo-EM: cryo-electron microscopy
RBD: RAS-binding domain
CRD: cysteine-rich domain
KD: kinase domain
CVs: collective variables.

## Supplementary Information for

**FIGURE S1.**
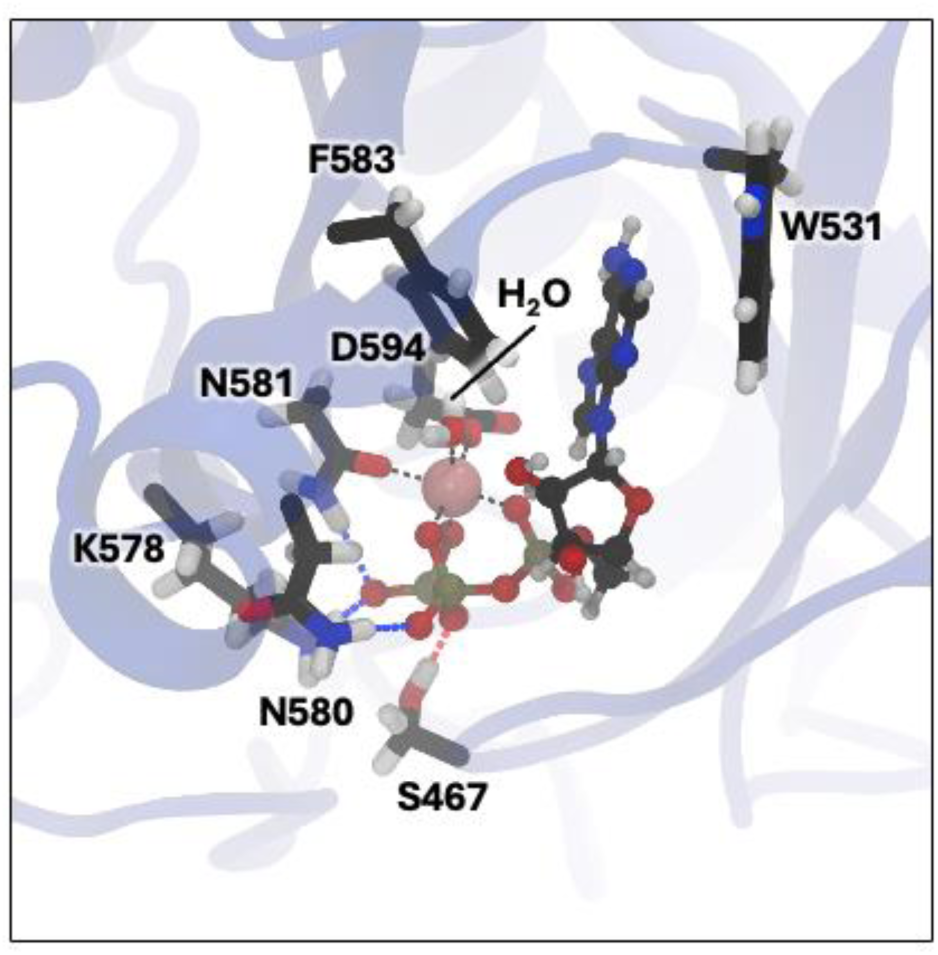
ATP binding site in KD. ATP and its coordinating Mg^2+^ ion were placed using MOE. ATP-Mg initial location and contacting amino acids were based on PDB 6U2G (chain B) followed by a short energy minimization to relax the ATP-Mg complex. The image shows the KD (blue), focused on the ATP binding pocket highlighting the ATP, Mg^2+^ (pink sphere) and contacting residues.

**FIGURE S2.**
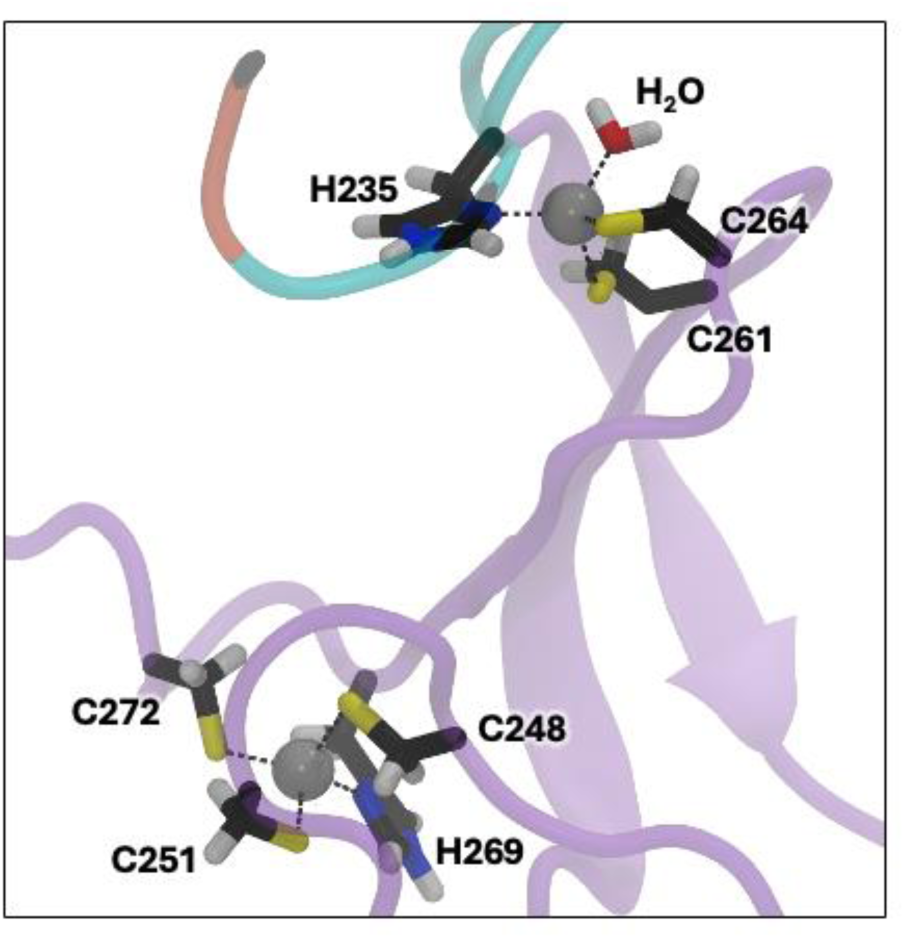
Zinc coordination sites in CRD. The reference structure PDB 7MFD does not resolve the fourth coordination of one of CRD’s zinc (top). We completed the coordination of that zinc by adding a coordinating water molecule and placed it to ensure proper coordination of residue H235. The image shows the CRD’s two zinc finger motifs highlighting the Zn^2+^ atoms and the contacting residues. The CRD is shown in purple, the C-terminal end of the RBD in orange, and the RBD-CRD linker in cyan. The two Zn^2+^ ions are shown in grey spheres.

**FIGURE S3.**
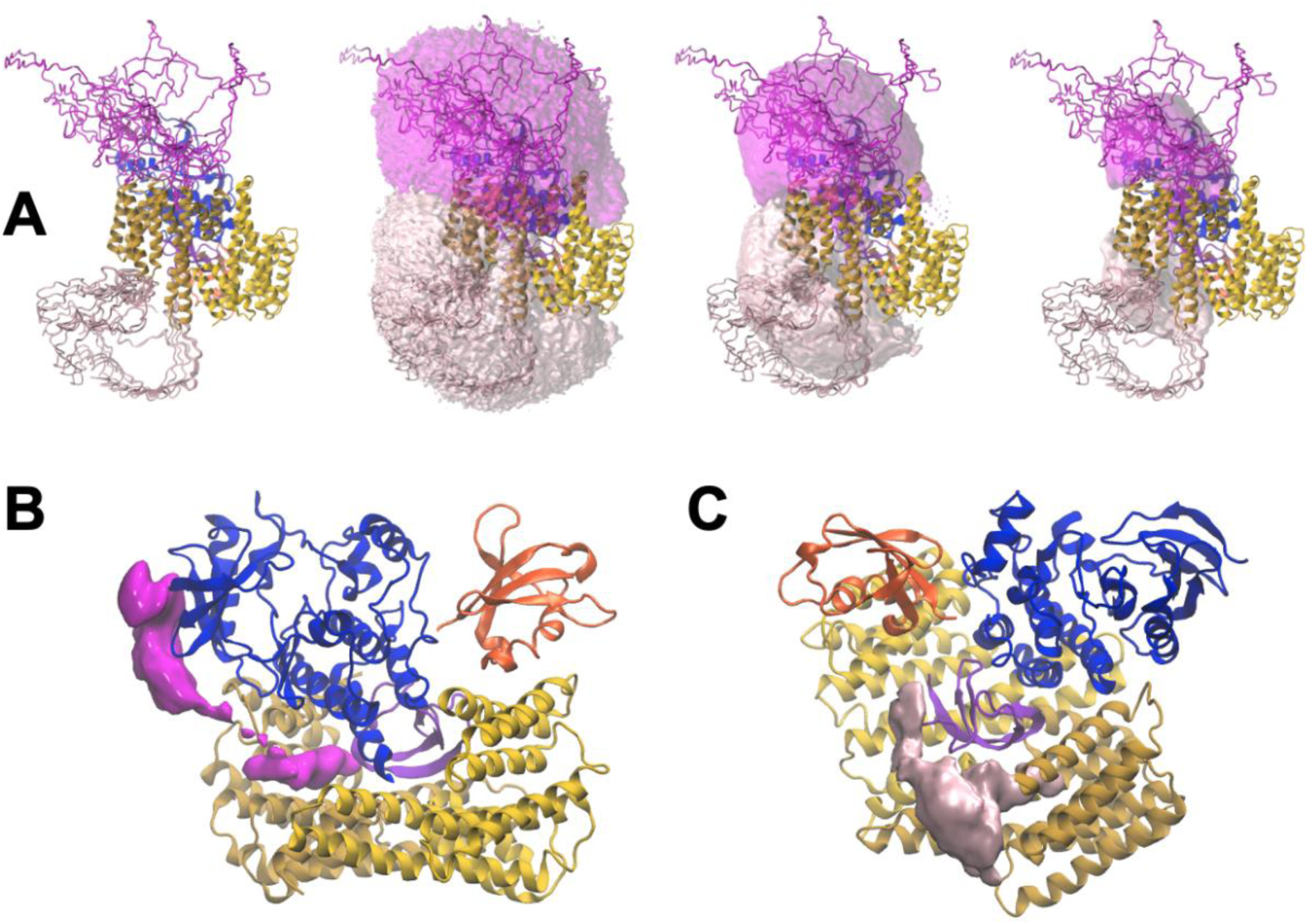
RAF loop placement and relaxation. Loops I (pink, residues 274 to 359) and II (magenta, residues 371 to 448) are disordered and are built into the model. The loops are built in with MOE and simulations are started with 27 different loop initial configurations, panel A. In the simulations the loops are very dynamic, sampling a larger confirmational space with the overall extent somewhat reduced compared to the initial placement, see example radius of gyration Fig. S4. A) The loops extended and common location are indicated with transparent loop density maps, showing low, medium, and high isovalues to illustrate the extended, commonly occupied, and frequently occupied areas for the loops. B) and (C) show zoomed in views of very high loop density areas for loop II and I, respectively, corresponding to where the loops are attached to the protein complex.

**FIGURE S4.**
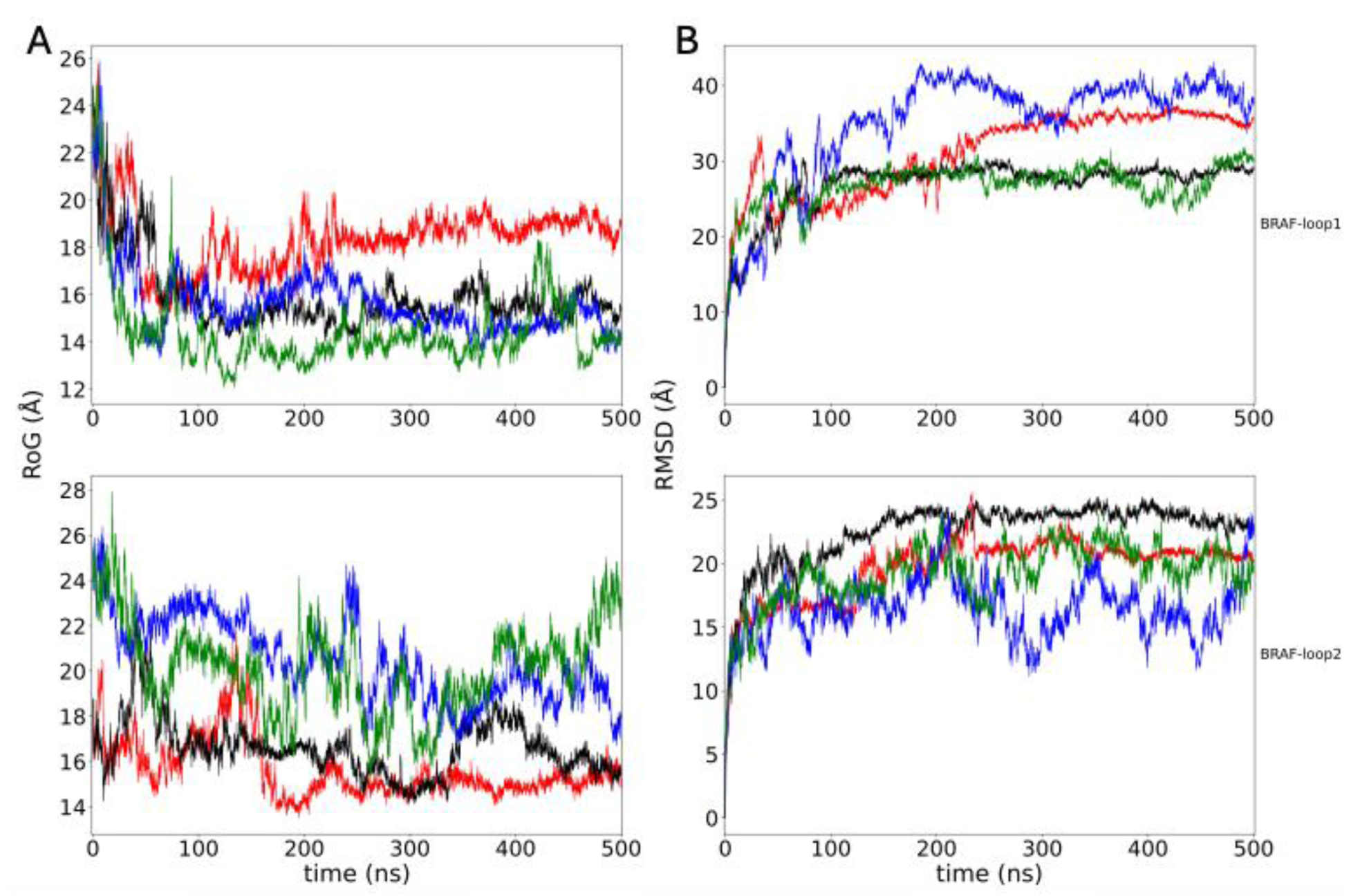
Extent and equilibration of the RAF loop configurations. Radius of gyration (RoG) (A) and RMSD (B) is shown for four random simulations of the 135 unbiased MD simulations. Top plots are for loop I (residues 282 to 359) and bottom plots for loop II (residues 371 to 448).

**Figure S5.**
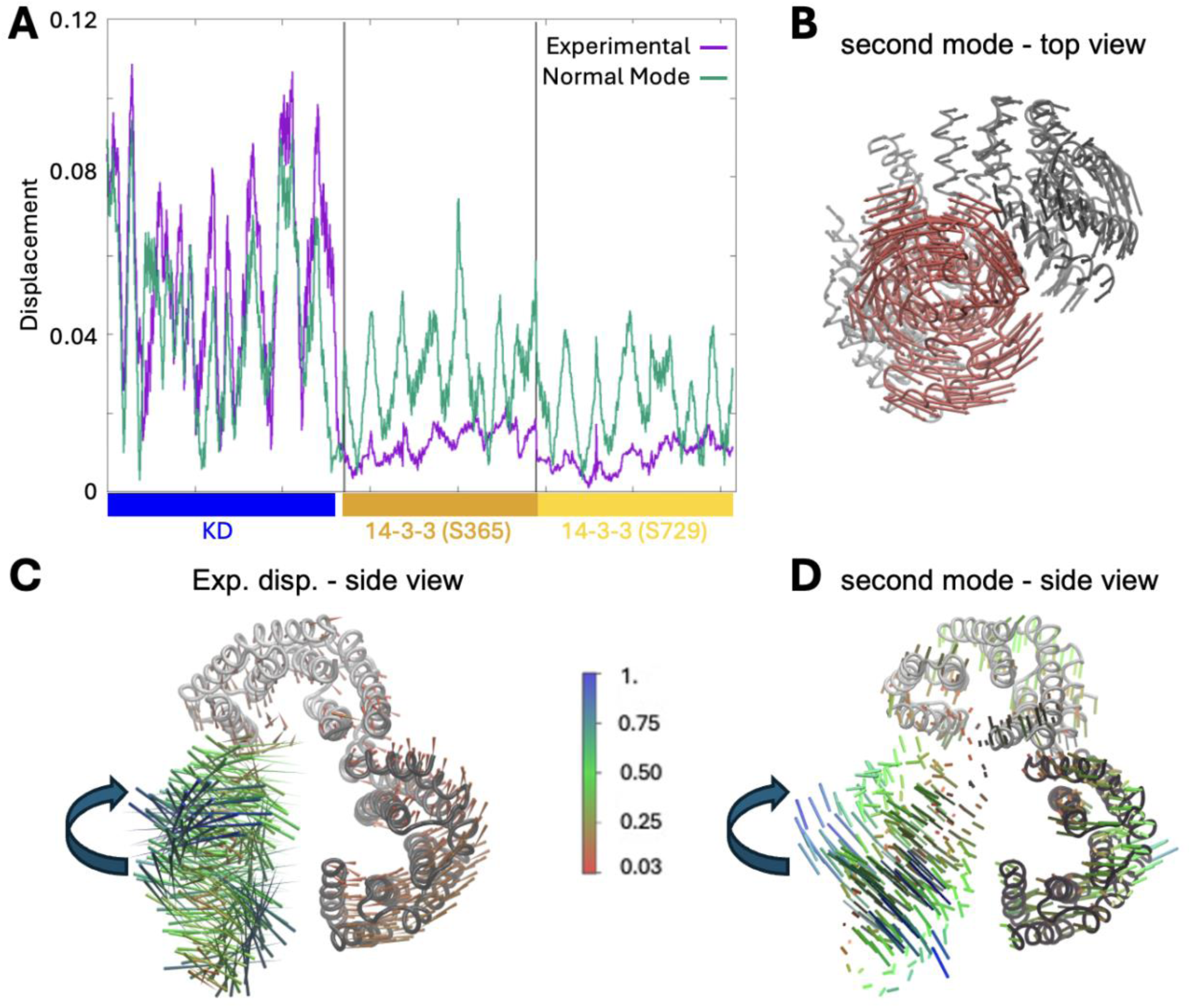
Details on the PCA analysis of the BRAF:14-3-3_2_ complex. A) Comparison of the best overlapping mode (second PCA mode) with experimental conformational displacement. B) Displacement according to computed second mode, top view. C) Displacement of 14-3-3_2_ and KD according to experimental conformational change PDB 7MFF to 7MFD. D) Displacement according to the computed second PCA mode. To assess whether the conformational flexibility around the autoinhibited state is related to the conformational changes required for the KD to transition between the monomeric and dimeric states, we compared these motions to the structural differences apparent between the dimerized BRAF (PDB: 7MFF) and monomeric BRAF (PDB: 7MFD) cryo-EM structures. We first superimposed the KD and 14-3-3 dimer from the monomeric BRAF structure 7MFD onto the KD and 14-3-3_2_ from the dimeric structure 7MFF. The RBD and CRD segments are discarded from 7MFD structure as well as from the PCA modes as these domains are not resolved in the dimer structure. Then we calculated the unit normalized displacement vector between 7MFD and 7MFF and compared the linear displacement required to transform the KD and 14-3-3_2_ complex. We show that the similarity of the vector is checked against the largest amplitude PCA modes. The second mode has a rotation type of movement for kinase. This second mode displayed highest similarity (dot product of the two vectors is −0.44) to the conformational change in between the cryo-EM structures. In panel A we show the experimental displacement vector together with the second PCA mode; 14-3-3_2_ domain is more mobile in the PCA mode, while the KD region displacements seem similar. In panel B the same PCA vector is projected onto the structure with arrows from top view of the rotation axis of the kinase. The rotational motion of kinase around the same axis is visible in experimental displacement in between 7MFD and 7MFF (panel C) and the second mode obtained from the PCA analysis of the MD trajectory (panel D).

**Table S1.** Details on the domains/residues in the BRAF:14-3-3_2_ model. Column 1 lists the names of each domain/region. Columns 2 and 3 list the starting and ending residue numbers for each region respectively. Column 4 describes whether the position of those residues were determined from the cryo-EM structure (PDB: 7MFD) or added to the structure by the modeling approaches described in the main text.

| Domain/region | Start residue | End residue | Cryo-EM or modeled? |
| --- | --- | --- | --- |
| RBD | 156 | 201 | Cryo-EM |
|  | 202 | 203 | *Modeled* |
|  | 204 | 227 | Cryo-EM |
| RBD-CRD linker | 228 | 234 | *Modeled* |
| CRD | 235 | 273 | Cryo-EM |
| CRD-zinc loop | 274 | 281 | *Modeled* |
| Loop I | 282 | 359 | *Modeled* |
| pS365 region | 360 | 370 | Cryo-EM |
| Loop II | 371 | 448 | *Modeled* |
|  | 449 | 456 | Cryo-EM |
| KD | 457 | 723 | Cryo-EM |
| PS729 region | 724 | 738 | Cryo-EM |
| 14-3-3 zeta (S365) | 2 | 69 | Cryo-EM |
|  | 70 | 72 | *Modeled* |
|  | 73 | 203 | Cryo-EM |
|  | 204 | 209 | *Modeled* |
|  | 210 | 230 | Cryo-EM |
| 14-3-3 zeta (S729) | 2 | 69 | Cryo-EM |
|  | 70 | 71 | *Modeled* |
|  | 72 | 133 | Cryo-EM |
|  | 134 | 136 | *Modeled* |
|  | 137 | 230 | Cryo-EM |

